# Defensive mutualisms affect plant and ant island biogeography at a global scale

**DOI:** 10.1101/2025.08.15.670609

**Authors:** Drin Brown, Megan E. Frederickson

## Abstract

**Aim:** Mutualism can shape biogeographic patterns at macroecological scales. Recently, Delavaux et al. (2024) found that plant mutualisms with mycorrhizae, nitrogen-fixing bacteria, and animal pollinators limit island colonization, weakening the latitudinal diversity gradient (LDG) on oceanic islands. Here, we revisit their analysis and examine whether ants and plants engaged in a common defensive mutualism mediated by extrafloral nectaries (EFN) are similarly under-represented on oceanic islands, and whether this defensive ant-plant mutualism also weakens the LDG on islands.

**Location:** Global.

**Time period:** Present-day species distributions and traits.

**Major taxa studied:** Vascular plants and ants.

**Methods:** We used global trait and occurrence databases to compare ant and plant species richness between oceanic islands and their likely source mainlands for taxa that do and do not interact mutualistically via EFNs. When analyzing the factors that determine whether a plant species occurs on oceanic islands, we also included the mutualism types studied by Delavaux et al. (2024), namely biotic pollination, mycorrhizal fungi, and nitrogen-fixing bacteria, and added plant habit, life history, and phylogeny as covariates.

**Results:** Both plants with EFNs and EFN-visiting ants are significantly over-represented on islands. The species richness of EFN-visiting ants and EFN-bearing plants also positively covary across islands. In ants, engaging in mutualisms mediated by EFNs significantly strengthens, rather than weakens, the LDG on islands, while for plants, EFNs have no effect or strengthen the LDG on islands, depending on data filtering. After accounting for plant habit, life history, and phylogeny, only biotic pollination significantly limits plant colonization of islands, whereas mycorrhizae and nitrogen-fixing bacteria now have positive or non-significant effects, respectively, on island colonization.

**Main conclusions:** EFNs facilitate rather than limit plant and ant colonization on islands. Mutualism does not ubiquitously limit island colonization, and some mutualism types strengthen rather than dampen the LDG on islands.

## Introduction

Reciprocally beneficial species interactions, or mutualisms, can shape biodiversity at macroecological scales. Mutualism can enlarge a species’ realized niche (Fowler et al., 2023) or geographic range (Afkhami et al., 2014, Nathan et al., 2023) and may be especially likely to facilitate range expansion if the interaction is generalized (Harrison et al., 2018), because local species may cooperate with the immigrating species and encourage its establishment (Chomicki et al., 2019). However, in specialized, obligate mutualisms, both partners may need to be present to establish in a new location, potentially limiting colonization opportunities for mutualist species (Simonsen et al., 2017, Delavaux et al., 2024, 2025). Here, we examined how defensive ant-plant mutualisms affect species richness on oceanic islands, where mutualism has been proposed to act as a strong filter on colonization (Delavaux et al., 2019, 2021, 2022, 2024).

Species richness on islands depends on the balance among colonization, speciation, and extinction or emigration, with island size and isolation from mainland sources of colonists being the primary determinants of species richness on islands (MacArthur and Wilson, 2001, Delavaux et al., 2021). However, biotic interactions may also limit colonization on islands, particularly on oceanic islands that were never part of a continent (Delavaux et al., 2024). Recently, Delavaux et al. (2019, 2021, 2022, 2024) found that mutualists have a reduced likelihood of colonizing oceanic islands, potentially because of a scarcity of compatible partners. Specifically, Delavaux et al. (2019, 2021, 2022, 2024) found that plants that associate with mycorrhizal fungi, nitrogen-fixing bacteria, and animal pollinators are more under-represented on islands than plants that lack these mutualistic associations, with important consequences for island ecosystems and biodiversity. For example, because biological nitrogen fixation is the primary means by which nitrogen enters the biosphere from the atmosphere, if the symbiosis between plants and N-fixing bacteria prevents these species from colonizing islands, island soils may not receive the same inputs of fixed N as mainland soils (Delavaux et al., 2022). Furthermore, although species richness generally increases from the poles to the equator, a pattern known as the latitudinal diversity gradient (LDG, Hillebrand, 2004; Kinlock et al., 2018), Delavaux et al. (2024) found that the mutualism filter on island species richness was strong enough to weaken the LDG on oceanic islands. In other words, because of mutualism, islands showed a different global pattern of biodiversity than mainlands (but see Pichler and Hartig, 2024). However, previous studies have focused only on nutritional and pollination mutualisms.

Here, we expand on the analysis of Delavaux et al. (2024) in several ways. Most importantly, we add trait data on another plant-animal mutualism: defensive ant-plant mutualisms mediated by extrafloral nectaries (EFNs). Over 4000 plant species produce EFNs, which are glands on plant parts other than inside flowers (Weber and Keeler, 2013). EFNs secrete sugar-rich nectar that attracts mainly ants and sometimes other arthropods that usually defend the plant from arthropod herbivores (Marazzi et al., 2013). Multiple meta-analyses across plant species, families, and biogeographic regions have shown that EFNs generally reduce herbivory and increase plant survival, growth, or reproductive success (Oliveira and Freitas 2004, Chamberlain and Holland 2009, Rosumek et al. 2009, Trager et al. 2010, Leal et al. 2023). Ant-plant interactions mediated by EFNs are thus a type of facultative and generalized mutualism—one in which the presence of the partner is usually not required for plant recruitment and partners are highly interchangeable (Lach et al., 2010, Marazzi et al., 2013). Globally, the richness of EFN-bearing plants follows the LDG (Luo et al., 2023). Phylogenetic studies have also shown that in general, the origination of EFNs in a plant lineage promotes diversification (Weber and Agrawal, 2014). This increase in diversity is expected because the presence of EFNs promotes population growth by providing defense, which can result in greater ecological opportunity leading to adaptive radiation. By the same token, EFNs may also accelerate plant range expansion (Blüthgen and Reifenrath 2003, Nathan et al., 2023). More EFN-bearing plants are legumes than any other plant family, and EFN-bearing legumes have spread to more new regions of the globe than legumes that do not have EFNs (Nathan et al. 2023). Thus, unlike the results of Delavaux et al. (2024) for plants that associate with mycorrhizae, N-fixing bacteria, and animal pollinators, we predicted that EFN-bearing plants may experience enhanced colonization on islands compared to plants that lack EFNs (Chomicki et al., 2019). If so, then counter to Delavaux et al. (2024), mutualism does not always filter species richness on islands, nor does it ubiquitously weaken the LDG on islands.

We also revisit the results of Delavaux et al. (2024) by accounting for plant phylogenetic relationships, growth form (i.e., habit, or whether a plant is woody or herbaceous), and life history, which were missing from their models of the species richness of mutualistic and non-mutualistic plants on islands. Mutualism does not evolve independently of other functional traits, making it challenging to disentangle the effects of mutualism from the effects of correlated traits on island biogeography. For example, Delavaux et al. (2019, 2021, 2024) examined how mycorrhizal fungi affect plant colonization on islands, but their models did not control for plant habit, which may be associated with both mycorrhizal status and insularity. Non-mycorrhizal plants are more often herbaceous, compared to ectomycorrhizal and arbuscular mycorrhizal plants, which are more often woody trees or shrubs (Soudzilovskaia et al., 2020) and herbaceous plants may be more likely than woody plants to colonize islands from mainlands (Schrader et al. 2020, 2025, see also Figure 1), although woodiness often evolves secondarily once herbaceous lineages establish on islands (Nürk et al. 2019, Zizka et al. 2022, Figure 1). Thus, plant habit, rather than mutualism *per se,* could explain the higher proportion of non-mycorrhizal plants on islands than mainlands (Delavaux et al. 2019). We sought to determine whether the finding of a mutualism filter on island plant diversity reported by Delavaux et al. (2019, 2021, 2022, 2024) persists after controlling for traits correlated with mutualism that also influence plant colonization on islands, especially plant habit and life history, and after accounting for the phylogenetic non-independence of plant species. Finally, although there are a growing number of studies of mutualism’s effects on global plant biogeography, few studies explore how mutualism shapes the global distribution of animals. We therefore also explored whether EFN-associated ants differ from non-EFN-associated ants in their propensity to colonize islands, and whether the richness of EFN-visiting ant taxa on islands is related to the richness of EFN-bearing plants. These analyses add significantly to our understanding of the factors that influence plant and ant diversity on oceanic islands.

**Figure 1.**
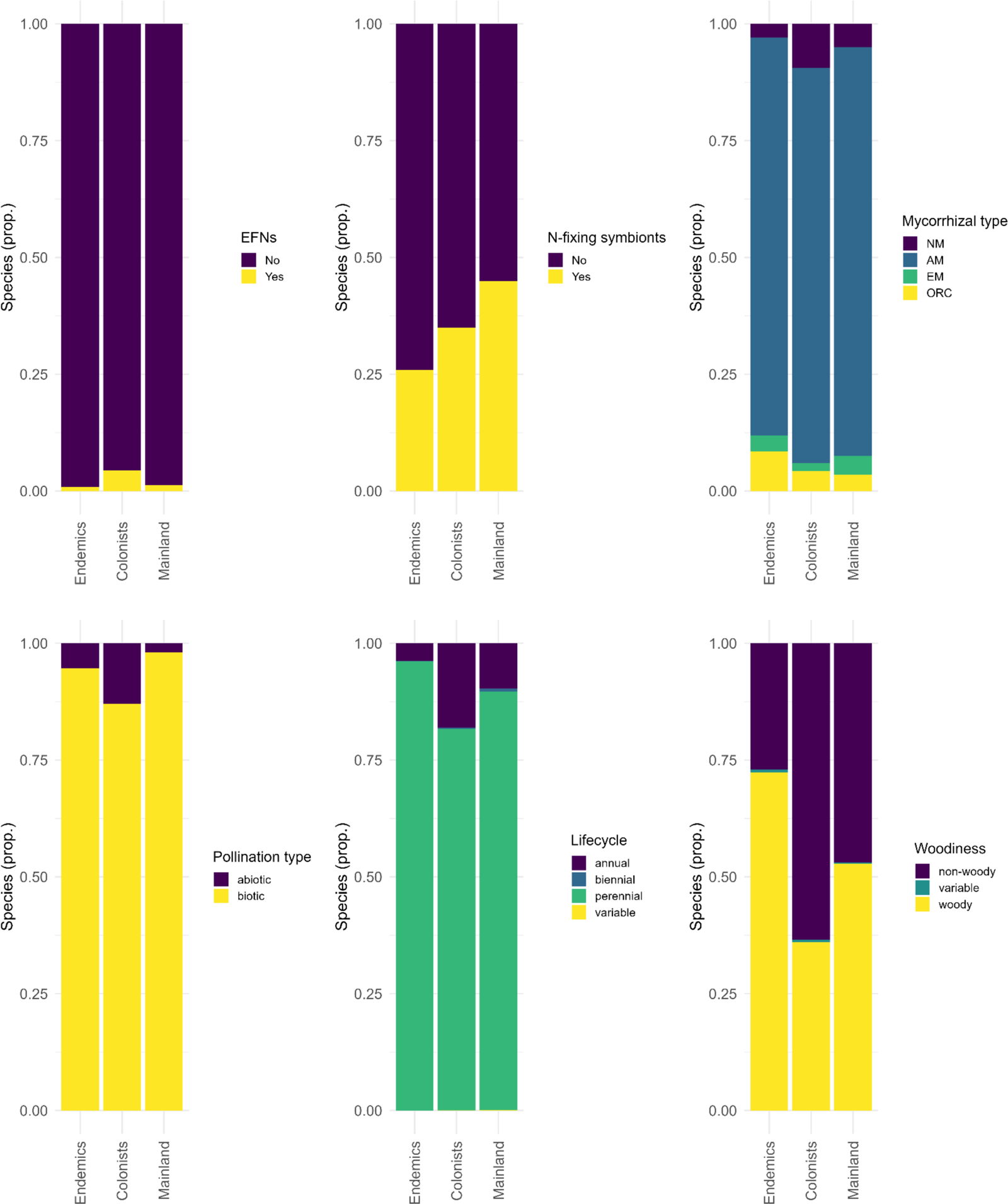
Plant traits among island endemics, island colonists, and mainland species after combining data from GIFT and Weber et al. (2024).

## Materials and methods

To examine the relationship between EFNs and plant colonization on islands, we used the Global Inventory of Floras and Traits (GIFT) database (Weigelt et al., 2020) for plant occurrence checklists, plant trait data (e.g., plant habit and life history), and geographic information and the World List of Plants with Extrafloral Nectaries (Weber et al., 2024) for plant EFN trait data. Figure 1 summarizes the raw plant trait data for species that occur on islands only (island endemics), species that occur on both islands and mainlands (colonists), and species that never occur on islands (mainland species). To determine whether visiting EFNs is associated with ant colonization on islands, we used the Global Ant Biodiversity Index (GABI) database (Guenard et al., 2017) and the Global Ant Biodiversity Index-Islands (GABI-I) database (Liu et al., 2023) for ant occurrence and geographic information and data from Kaur et al. (2019) and Novais et al. (2025) on which ant species visit EFNs. These data sources are also described in more detail in Table 1 and in the supplemental appendix. All analyses were performed in R Studio using R version 4.4.2 (R Studio R Core Team, 2024).

**Table 1.**
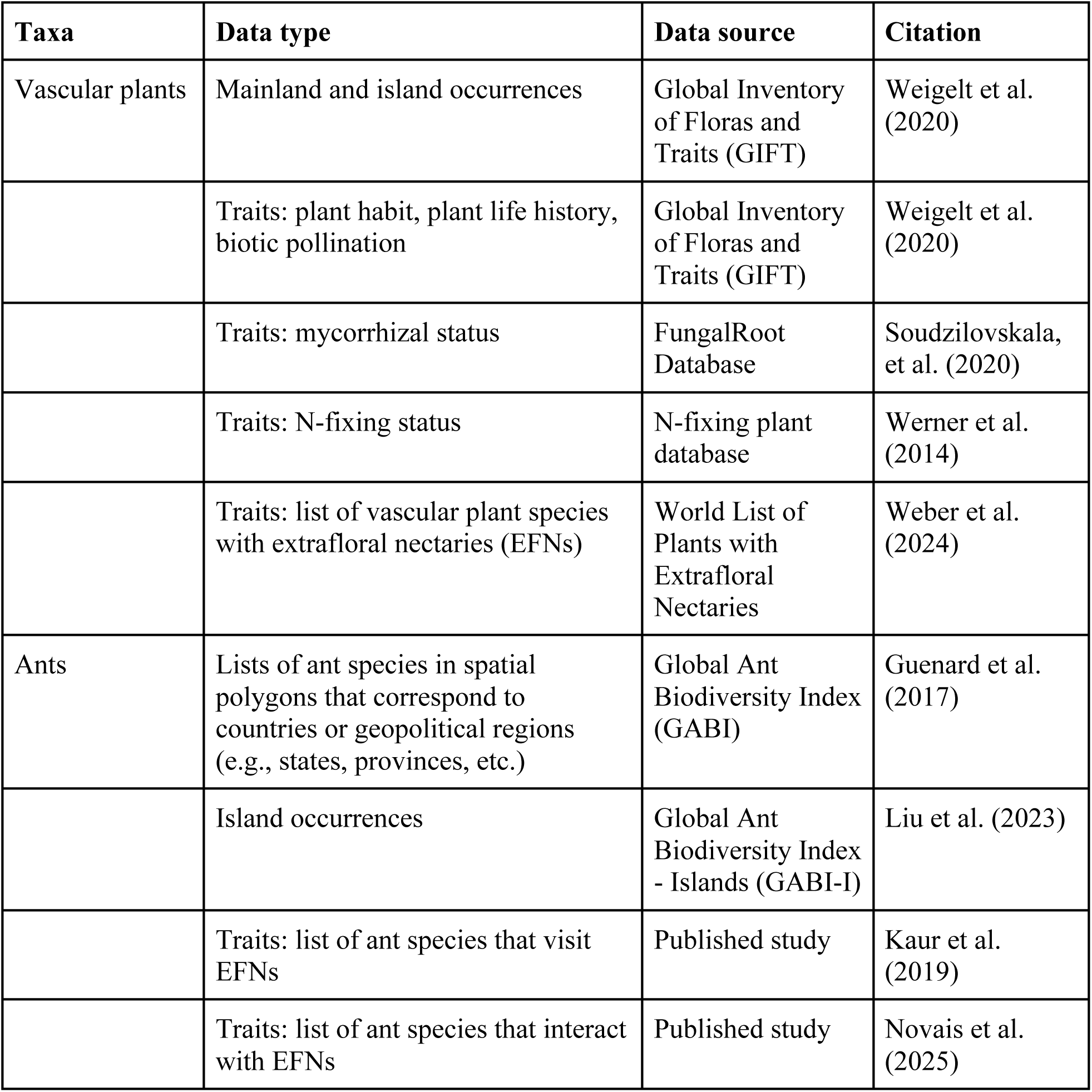
Summary of data sources and citations for geographic occurrence and trait data.

We fit two types of models to both the ant and the plant data: 1) species-level models in which each data point is a species, and the (binary) response variable is whether it occurs on islands, and 2) island-level models in which each data point is an island, and the (continuous) response variable is the log response ratio of island species richness to mainland species richness. We modelled the data two ways because the species-level models allowed us to account for phylogeny and, for plants, other traits like habit and life history, but not island characteristics, while the island-level models allowed us to account for the classical predictors of species richness on islands like island size and isolation, but not phylogeny or species-level traits. Previous work exploring the factors affecting island biodiversity has taken both approaches; for example, Zell et al. (2024) modeled island presence/absence as a binary response variable and a suite of species-level functional traits as predictors, while Delavaux et al. (2024) modelled what they termed the ‘island species deficit’ (1 the ratio of island to predicted mainland species richness), which is very similar to the log(island species richness/mainland species richness) metric we calculate here. We calculated log response ratios because of their well understood statistical properties (Hedges et al. 1999).

For the species-level plant model, we used data from Delavaux et al. (2024), which included a list of oceanic islands, vascular plant checklists, and trait data on associations with animal pollinators, N-fixing bacteria, and mycorrhizae. We added EFN trait status from the World List of Plants with Extrafloral Nectaries (Weber et al., 2024). We assumed that plants not in the Weber et al. (2024) list did not have EFNs, as this list is the most complete in the literature to date. As a result, we assigned an EFN trait status to all plant species in GIFT, with the result that 1.5% of the plant species in GIFT have EFNs (Figure 1). For comparison, Weber and Keeler (2013) estimate that EFNs occur in 2-3.6% of all flowering plant species.

In the species-level models, we treated island presence as a binary trait, such that each species either occurs on one or more oceanic islands (1) or does not occur on any oceanic island (0) and predictors estimate trait effects on island presence/absence. To account for the phylogenetic non-independence of species, we reconstructed a phylogeny of all vascular plants in the GIFT checklists using the R package V.Phylomaker2 (Jin & Qian, 2022) and then fit a phylogenetically adjusted GLM with island presence/absence as the response variable and EFN status, mycorrhizal association, nitrogen fixation, biotic pollination, plant habit, and plant life history as predictors. We also fit a species-level model for ants examining island presence/absence as a function of EFN association, using the ant phylogeny from Nelsen et al. (2018). For both species-level models, we used the phylogenetically adjusted logistic regression method of Ives and Garland (2010) as implemented in the phyloglm package in R.

For the island-level models, we needed to compare species richness between oceanic islands and mainland sources of colonizing species. To obtain the set of oceanic islands, we used data in Delavaux et al. (2024), which used the GIFT R package (Denelle et al., 2023) to extract geographic data from the GIFT database, including information on island geological origins, which distinguishes oceanic from non-oceanic islands. Following Delavaux et al. (2024), we analyzed only oceanic islands because they were not originally part of a mainland, so they represent unique archives of evolution and colonization. To have a measure of mainland species richness against which to compare island species richness, Delavaux et al. (2024) interpolated mainland species richness using only absolute latitude as a predictor. Pichler and Hartig (2024) re-analysed the same data using random forest models parameterized with either latitude alone or latitude and longitude to predict mainland species richness; however, they found no support for mutualism dampening the LDG on oceanic islands with these models (but see Delavaux et al., 2025). Instead of using a model to predict mainland species richness, we compared species richness on oceanic islands with the observed species richness of their five most likely source mainlands, calculated as the total number of unique species across the five mainlands (i.e., if a species occurred in more than one of the five source mainlands, it was only counted once). To identify the five most likely source mainlands for each island, we adopted the approach of König et al. (2021). König et al. (2021) modelled species turnover as a function of both environmental and geographic distance between islands and mainlands and then minimized species turnover to predict the most likely source mainlands for each island in GIFT. For plants, we constrained the possible source mainlands to those with vascular plant checklists available in GIFT, and for ants, we use GABI geographic polygons as possible source mainlands. To model ant species turnover between islands and mainlands and the correlation between ant and plant species richness on islands (see below), we needed to associate the ant data from GABI and GABI-I with the GIFT geographic entities and their environmental metadata. We spatially joined ant occurrence data from GABI and GABI-I to the geographic data in GIFT by latitude/longitude. Ant occurrence data in GABI-I had latitudes and longitudes that sometimes differed from the coordinates supplied for islands in GIFT. We used the sf package (Pebesma, 2023) in R to generate a distance matrix between each island as represented in GIFT and in GABI-I, using the minimum distance to pair GABI-I and GIFT islands, and then used a 1km distance cutoff for a pair to be treated as the same island.

In order to exclude understudied species in which the presence or especially the absence of a mutualistic trait might be unreliable (Chamberlain et al., 2022), for both plants and ants, we used species occurrence records from the Global Biodiversity Information Facility (GBIF, package rgbif, Chamberlain et al., 2022) to filter out species that are very rare (< 100 GBIF occurrences). We present results with this filter applied below, but there were almost no differences in statistical significance or direction of effect in models with versus without this filter, except for the interaction effect between latitude and EFN status in the island-level plant model that was non-significant with the filter (Table 3) but statistically significant without it (Table S3).

When testing for a mutualism filter on island colonization (as in Delavaux et al., 2024), the most relevant species on islands are those present in their likely source mainlands (Konig et al., 2021). As such, we required that for an island species to be counted, it had to be present in at least one of that island’s five most likely source mainlands. This excluded island endemics, which likely originated via speciation on islands and not as colonists from mainlands, as well as island species that arrived from mainlands not captured in our source pool for each island; for comparison, we also include models with all island species in the supplemental material (Table S1 and Figures S1 and S2).

For each island, we calculated the log response ratio (LRR) of island to source mainland species richness for plants with and without EFNs and ants that do or not visit EFNs and then fit generalized linear models (GLMs) separately for the ant and plant trait data. Because log response ratios can be calculated only from positive, non-zero values, we added a constant of 1 to all count data. The main predictor of interest was whether plants have EFNs, or ants visit EFNs. Additional biogeographical and climatic variables in the models included absolute latitude of the island in each island-mainland group, average distance from island to source mainlands, island area and elevational range, average mainland area and elevational range, and mean annual precipitation on the island. We log-transformed all geographic variables to improve normality of the residuals.

Finally, to examine the relationship between plant and ant species richness, we also fit separate GLMs with the island species richness of either EFN-visiting or non-EFN-visiting ants as response variables and the species richness of EFN-bearing and non-EFN bearing plants, as well as biogeographical variables, as predictors. These models tested whether the diversity of ant defenders is correlated with the diversity of plants with EFNs, which would be expected if ant diversity begets plant diversity, or *vice versa,* thanks to ant-plant mutualisms mediated by EFNs..

## Results

Visualizing the raw trait data shows that among the 77,874 plant species in GIFT with appropriate trait data, island colonists (species that occur on both islands and mainlands) are more likely to be pollinated by abiotic agents (especially wind), to be non-mycorrhizal, and to lack nitrogen-fixing symbionts than mainland-only plant species, recapitulating the general finding of Delavaux et al. (2024) that these plant mutualisms are less well represented on oceanic islands than on mainlands (Figure 1). However, island colonists are more likely, not less likely, to have EFNs (Figure 1). Interestingly, island endemics are more likely to be biotically pollinated, mycorrhizal plants that lack EFNs and nitrogen-fixing symbionts than island colonists (Figure 1).

Figure 1 also reveals that island colonists are more often herbaceous annuals than either island endemics or mainland-only plant species, meaning it is important to control for plant habit and life history when modelling the influence of plant mutualisms on island colonization. We fit a species-level model to the plant data that included these traits and the four mutualism-related plant traits, and confirmed that, indeed, plants on oceanic islands are significantly more likely to be herbaceous annuals, compared to woody or perennial plants (Table 2). In this model, EFN-bearing plants were also significantly more likely to occur on oceanic islands than plants without EFNs (Table 2), as were plants with arbuscular mycorrhizal fungi; for AMF, this is the opposite of what Delavaux et al. (2024) found, but they did not account for plant habit and life history. Nonetheless, consistent with Delavaux et al. (2024), biotically pollinated plants were significantly under-represented on islands, compared to abiotically pollinated plants (Table 2, Figure 2). There was no significant effect of nitrogen-fixing symbionts on island presence (Table 2, Figure 2). In the ant species-level analysis, EFN-visiting ants were also significantly more likely to occur on oceanic islands than ant species that do not visit EFNs (Table 2). There was also phylogenetic signal for whether ant and plant species occur on islands, with ants having stronger phylogenetic signal for insularity than plants (see Table 2 for alphas).

**Figure 2.**
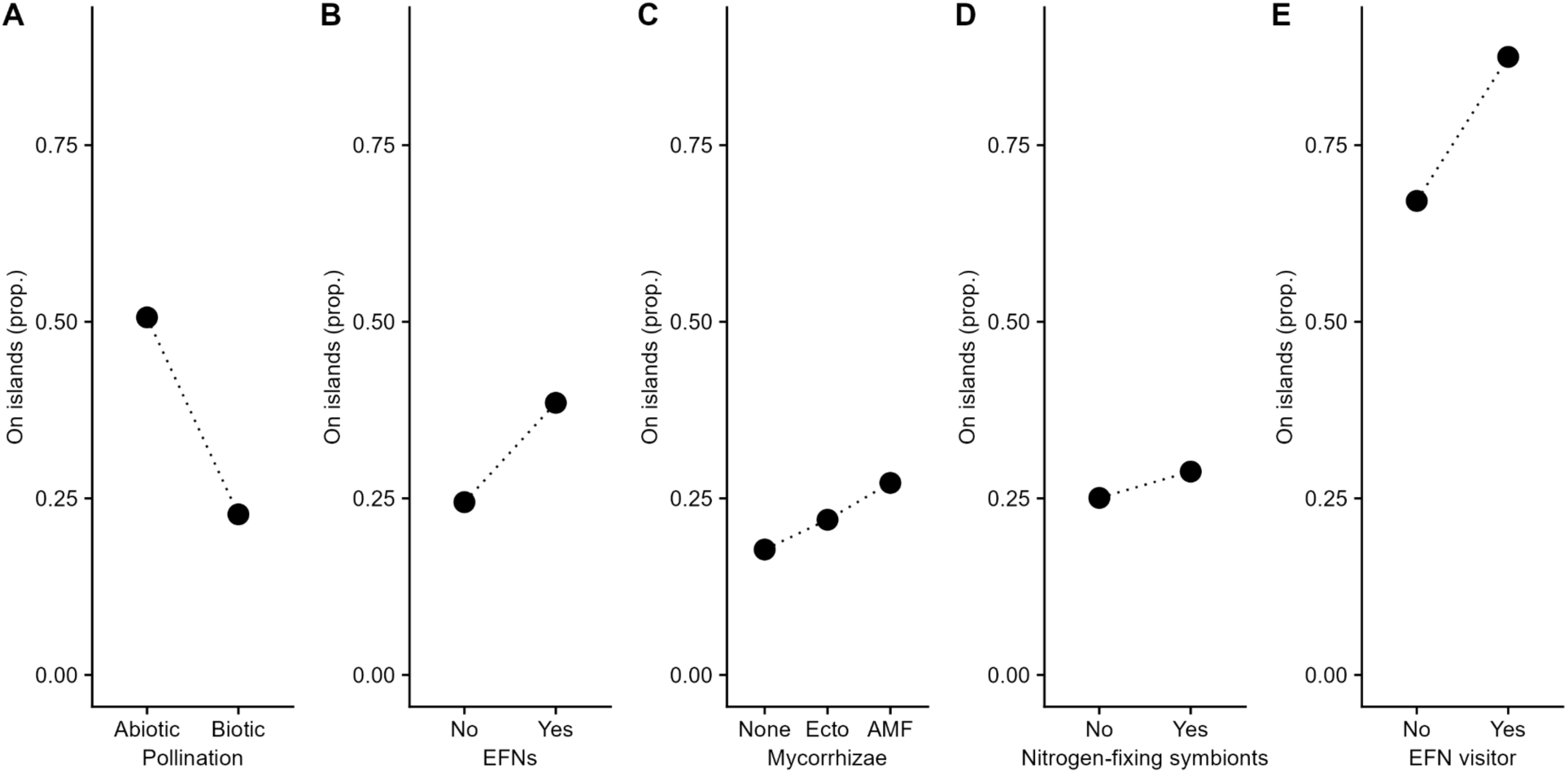
Proportions of plant (A-D) or ant (E) species that occur on islands, for plant species that A) are biotically versus abiotically pollinated, B) do or do not have EFNs, C) associate with arbuscular mycorrhizal fungi (AMF), ectomycorrhizal fungi (ecto), or are non-mycorrhizal (none), and D) do or do not associate with nitrogen-fixing symbionts, and for ant species that E) do or not visit EFNs.

**Table 2.** Estimates and standard errors (s.e.) from phylogenetic logistic regression models for A) plants (N = 1517 species with complete trait data) and B) ants (N=1660 species with trait data). Island species presence/absence was modelled as a function of EFN status only for ants, or EFN status, mycorrhizal association, nitrogen-fixing symbionts, biotic pollination, woodiness, and lifecycle for plants. Alpha is a measure of phylogenetic signal. In A), the intercept represents abiotically pollinated, annual, herbaceous plants with no EFNs, no mycorrhizae, and no nitrogen-fixing symbionts. In B) the intercept represents ants that do not visit EFNs.

|  | Estimate | s.e. |
| --- | --- | --- |
| **A - Plants** |  |  |
| Intercept | 0.79 | 0.42 |
| *EFN* | 0.91*** | 0.20 |
| *Arbuscular mycorrhizal* | 0.68* | 0.29 |
| Ectomycorrhizal | 0.32 | 0.38 |
| Orchid mycorrhizal | 1.18 | 1.45 |
| Nitrogen fixer | 0.19 | 0.23 |
| *Biotic pollination* | -1.19*** | 0.24 |
| Variable woodiness | -13.07 | 840.39 |
| *Woody* | -0.96*** | 0.19 |
| Biennial | -1.06 | 1.06 |
| *Perennial* | -0.69* | 0.30 |
| Variable lifecycle | -13.64 | 793.66 |
| Mean tip height | 137.69 |  |
| Alpha | 0.046 |  |
| B - Ants |  |  |
| *Intercept* | *0.664\*\*\** | *0.078* |
| *EFN status* | *1.070\*\*\** | *0.189* |
| Mean tip height | 136.99 |  |
| Alpha | 0.055 |  |

Significant effects are *italicized* and indicated by \*\*\* for $p \leq 0.001$ , \*\* for $p \leq 0.01$ and \* for $p \leq 0.05$ .

The mainland-island grouping procedure yielded 169 groups (of one oceanic island and five likely source mainlands) for plants (Figure 3A) and 178 for ants (Figure 4A). The number of mainland-island groups differed between plants and ants because of differences in data coverage. Log response ratios of island to mainland species richness were always negative, reflecting the well-known deficit of species on islands relative to mainlands (Carlquist, 1974). In the island-level plant model, EFN-bearing plants were again over-represented on oceanic islands, meaning they were more likely to occur on oceanic islands than plant species without EFNs (Table 3A, Figure 3B); the LRR of island to mainland species richness was less negative for plants with than without EFNs, indicating a smaller island species deficit for plants with EFNs. For plants, islands that were farther from their source mainlands, smaller islands, islands with smaller source mainlands, and drier islands all had larger species deficits compared to mainlands (i.e., more negative LRRs of island to mainland species richness) (Table 3A).

**Figure 3.**
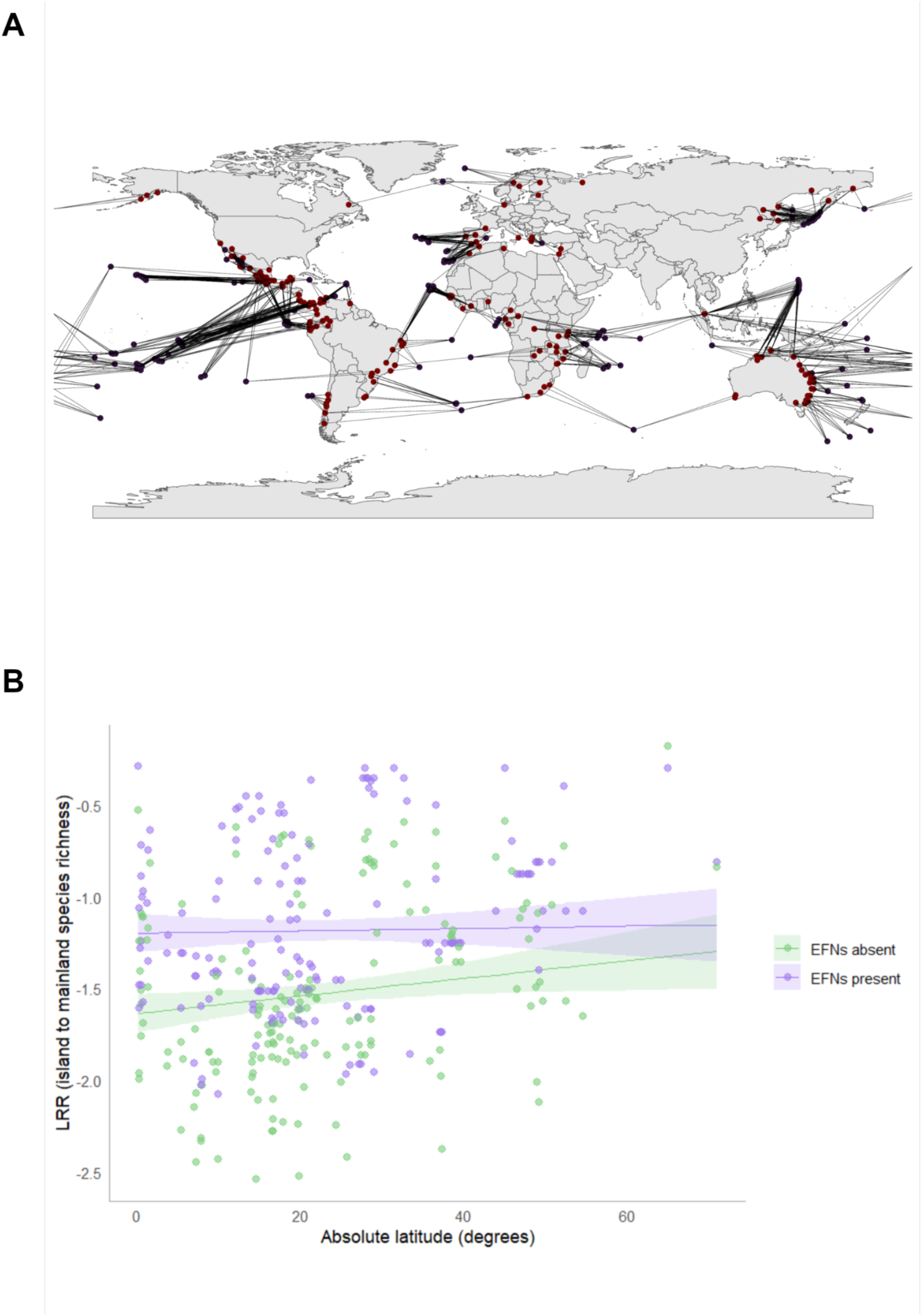
A) Map of five likely source mainlands (red) for each island (blue) for plants. B) Plot of log response ratio of island to mainland species richness for EFN- (purple) and non-EFN-bearing (green) plants against absolute latitude of the island.

**Figure 4.**
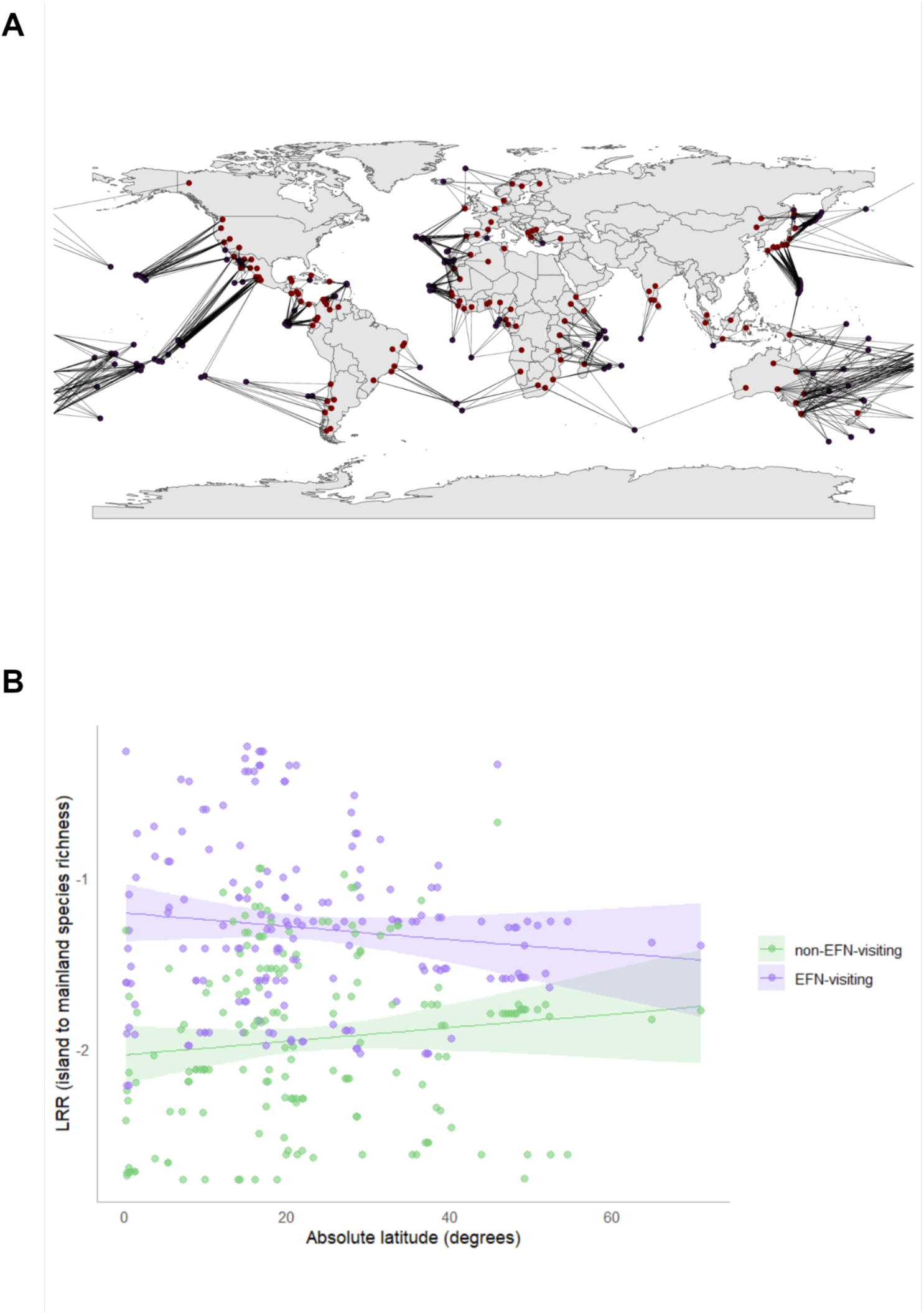
A) Map of five likely source mainlands (red) for each island (blue) for ants. B) Plot of log response ratio of island to mainland species richness for EFN-visiting (purple) and non-EFN-visiting (green) ants against absolute latitude of the island.

**Table 3.** Estimates and standard errors (s.e.) from Gaussian generalized linear models of the log response ratio of island to mainland species richness for A) plants (N = 169 groups) and B) ants (N = 178 groups) as a function of EFN mutualism status and biogeographical variables. Only island species that occur in at least one of the five likely source mainlands per island were included. Parameters ending in .i are island, .ml are mainland. Here, a positive estimate means a reduction of the island species deficit, whereas a negative estimate means a strengthening of the deficit.

|  | Estimate | s.e. |
| --- | --- | --- |
| **A - Plants** |  |  |
| *Intercept* | -1.64*** | 0.05 |
| *Absolute latitude.i* | 0.005* | 0.002 |
| *EFN?* | 0.44*** | 0.07 |
| *Distance.ml* | -0.23*** | 0.03 |
| *Area.i* | 0.22*** | 0.03 |
| *Area.ml* | 0.17*** | 0.05 |
| Elevational range.i | 0.004 | 0.03 |
| Elevational range.ml | 0.05 | 0.03 |
| *Annual precip.i* | 0.19*** | 0.03 |
| Latitude * EFN | -0.004 | 0.003 |
| **B - Ants** |  |  |
| *Intercept* | -2.032*** | 0.086 |
| Absolute latitude.i | 0.004 | 0.003 |
| *EFN?* | 0.829*** | 0.092 |
| *Distance.ml* | -0.072* | 0.035 |
| *Area.i* | 0.091** | 0.026 |
| Area.ml | -0.038 | 0.051 |
| Annual precip.i | -0.046 | 0.034 |
| *Latitude \* EFN* | -0.008* | 0.003 |

Significant effects are *italicized* and indicated by \*\*\* for $p \leq 0.001$ , \*\* for $p \leq 0.01$ and \* for $p \leq 0.05$ .

In the island-level plant model, there was also a significantly positive main effect of latitude, meaning that tropical islands are more plant species-poor relative to tropical mainlands than are temperate islands relative to temperate mainlands (Table 3A, Figure 3B). In this model, in which we filtered the data to exclude very rare species for which trait data may be unreliable, there was no significant interaction between latitude and EFN status, indicating that the effect of EFNs on island plant species richness does not vary with latitude (Table 3A, Figure 3B). In contrast, when we did not filter out rare plant species using a minimum cutoff of 100 GBIF occurrences, there was a statistically significant interaction between latitude and EFN status, such that the effect of EFNs on the LRR of island to mainland species richness is most positive at the equator and less positive at high latitudes (i.e., a negative EFN x latitude effect on the LRR, Table S3A, Figure 3B). Thus, in the unfiltered data, EFNs increase island plant species richness more near the equator than near the poles, and thereby strengthen the LDG on oceanic islands. In other words, EFNs either have no effect or strengthen the LDG for plant species on oceanic islands, depending on how the data are filtered.

In the island-level ant model, EFN-visiting ants were also over-represented on islands (Table 3B, Figure 4B); the LRR of island to mainland species richness was less negative for ants that visit EFNs than for ants that do not visit EFNs. Island area was also a significant predictor in the island-level ant model, again with smaller islands having a larger species deficit, as evidenced by a more negative LRR of island to mainland species richness (Table 3B). For ants, there was a significant negative interaction between island latitude and EFN visitation, in both the filtered (Table 3B) and unfiltered (Table S3B) data. Thus, regardless of how we filtered the ant data, visiting EFNs strengthened the LDG for ant species on oceanic islands (Table 3B, Figure 4B).

Because species that occur only on islands, which we assume to be endemics, differ from island colonists in mutualistic traits (Figure 1), in Table 3, we modelled the log response ratio of island to mainland species richness including only island species that also occur on a corresponding mainland and likely arrived from there as colonists. When we included all island species, adding island endemics and island species originating from beyond the most likely source mainlands, a significantly positive effect of EFN trait status remained for ants, but not plants (Table S1, Figures S1 and S2). In these models, EFN trait status never interacted significantly with latitude, for either ants or plants (Table S1). Thus, including island endemics, which have different traits than colonists (Figure 1), obscures the relationships among EFNs, island colonization, and latitude that emerge when we limit our analyses to species that colonize islands from mainlands (as in Table 3).

Across oceanic islands, the species richness of island ants that visit EFNs was correlated with the species richness of plants with EFNs, but not with the species richness of plants that do not have EFNs (Table S2A, Figure S3). In contrast, the species richness of ants that do not visit EFNs was significantly correlated with the species richness of non-EFN-bearing plants, but not with the richness of plants with EFNs (Table S2B, Figure S3). The richness of EFN-visiting ants, but not other ants, on islands followed the typical latitudinal diversity gradient (e.g. Willig, 2003), with more species in the tropics than at high latitudes, and there were also more species of EFN-visiting ants on larger islands and islands nearer to mainlands (Table S2A). The richness of ant species that do not visit EFNs was significantly predicted by only the richness of non-EFN-bearing plants and by annual precipitation, with wetter islands having more non-EFN-visiting ant species (Table S2B).

## Discussion

Ant-plant mutualisms mediated by EFNs are generalized and facultative (Marazzi et al., 2013), and previous work has shown that this mutualism facilitates plant and ant introductions to new ranges (Nathan et al., 2023) and spurs plant diversification (Weber and Agrawal, 2014). As such, we expected that, contrary to Delavaux et al. (2024), this mutualism type would not act as a filter on the diversity of colonizing species on oceanic islands, and might instead enhance island colonization from mainlands. The results of both the species- and island-level models support our prediction; plants that have EFNs and ants that visit EFNs are over-represented on oceanic islands, compared to taxa lacking these traits. Furthermore, in the island-level models, the positive effect of this ant-plant mutualism on island colonization either does not vary latitudinally (Table 3A) or increases towards the equator (Table 3B, Table S3), meaning that engaging in mutualism mediated by EFNs either has no effect or intensifies, but never weakens, the LDG on islands. Nathan et al. (2025) found that the castor bean, *Ricinus communis*, invested more in EFNs in lower than higher latitudes, and that more ants recruited to these EFNs nearer the equator. Our results find support for this pattern of higher representation of ant-plant mutualisms mediated by EFNs at lower latitudes globally, in agreement with the Biotic Interactions Hypothesis (BIH) (Schemske et al., 2009). Finally, the richness of EFN-visiting ants and EFN-bearing plants positively covary across oceanic islands, suggesting that the mutualistic ant and plant taxa that interact via EFNs either respond in similar ways to climatic or other environmental variation among islands, or are subject to positive feedback fueled by the mutualism itself.

The raw proportions of mutualistic versus non-mutualistic taxa vary among island endemics, colonists, and mainland plant species in ways that are consistent with Delavaux et. al (2021, 2022, 2024); colonists are less likely than mainland plant species to have animal pollinators, mycorrhizae, and rhizobia (Figure 1). However, colonists are more likely than mainland species to have EFNs (Figure 1), suggesting that not all mutualism types affect island colonization equally, as further supported by the species- and island-level models. In contrast to the study by Delavaux et al. (2024), which fit only island-level models that did not account for other plant traits beyond mutualism, we fit a species-level model that included not only mutualism-related traits, but also plant habit and life history as covariates. Unsurprisingly, these covariates significantly predicted insularity; herbaceous, annual plants are over-represented on islands, compared to woody, perennial plants (Figure 1, Table 2). After controlling for plant habit and life history, biotic pollination still had a negative effect on whether plant species live on islands, as in Delavaux et al. (2024), but the effect of nitrogen-fixing symbionts was non-significant, and the effects of arbuscular mycorrhizal fungi and EFNs were significantly positive (Table 2, Figure 2); plants with EFNs and arbuscular mycorrhizal fungi are significantly more likely, not less likely to occur on on oceanic islands than non-EFN and non-mycorrhizal plant species. These findings cast doubt on the generality of the mutualism filter on island plant diversity reported by Delavaux et al. (2024).

In the species-level model, biotic pollination still significantly reduced the likelihood that a plant species occurs on islands, compared to abiotic pollination, and indeed Delavaux et al. (2024) found that biotic pollination results in the largest island species deficit among the mutualism types they examined. Delavaux et al. (2024) ascribe this mutualism filter on island colonization to “the reduced likelihood of simultaneous co-colonization of partners” on islands. However, this mechanism should mainly affect plants that are highly specialized on a specific taxon of animal pollinator, whereas most plant-pollinator interactions are highly generalized and dynamic (Waser et al., 1996). Some mutualisms can also re-assemble even if colonization by the partners is not simultaneous (Hembry and Tocora, 2025). Nonetheless, the idea that animal pollination is less reliable on islands than on mainlands is old and has solid empirical support; both selfing and wind pollination can be favored on islands (Barrett 1996, Grossenbacher et al. 2017). Wind pollination may be favored not only because of a dearth of animal pollinators, but also because open, wind-swept coastal habitats on islands favor plant groups, like grasses, in which wind pollination is widespread (Barrett et al. 1996, Friedman and Barrett, 2009).

Delavaux et al. (2024) did not distinguish between taxa that arrived on islands from elsewhere and taxa that have radiated on islands–i.e., whether a plant species has allochthonous or autochthonous origins. For the island-level models, we excluded potentially autochthonous island endemics because they do not give information about the limits to colonization; instead they may reflect the evolutionary drivers on the island itself, although endemics still evolve from successful colonists, and so may retain their traits. Comparing the traits of island endemics and colonists, as in Figure 1, reveals differences in their mutualism-related traits; mutualism may affect the process of dispersing to and establishing on an island differently from how mutualistic traits and taxa subsequently evolve on islands. For example, while reduced dependence on animal pollinators might favor island colonization, adaptive radiation and diversification on islands might depend on ‘escaping homozygosity’ by outcrossing (Barrett 1996); this is consistent with the greater proportion of biotically pollinated plants among island endemics than island colonists in Figure 1. Similarly, although plant species arriving on islands from mainlands are more likely to have EFNs than plants that do not colonize islands (Tables 2A and 3A), island endemics are less likely to have EFNs than colonists or mainland taxa (Figure 1), consistent with previous work that has found that some island endemics have lost EFNs (Sugiura et al. 2026), perhaps because of reduced herbivory or ant abundances on some oceanic islands (e.g., in Hawaii, Keeler 1985). When we observe a species living on islands as having particular traits, it reflects the end result of multiple processes, including dispersal, successful establishment in the face of abiotic factors and biotic interactions, and subsequent adaptation and speciation on islands (Losos and Ricklefs, 2009), that are difficult to capture in coarse comparisons of the traits of island and mainland taxa. Mutualism is likely to impact many of these processes, shaping island faunas and floras.

We also added a new mutualism type to the Delavaux et al. (2024) analysis: the mutualism between plants with EFNs and their ant defenders. Although this mutualism type is much rarer among plants than biotic pollination (Figure 1), these two mutualisms have similar effect sizes on insularity (Table 2, coefficient for biotic pollination is -1.19, compared to +0.91 for EFNs), meaning that they have almost equivalent effects on shaping island floras, albeit in opposite directions. In other words, the coefficients of the species-level model indicate that a biotically pollinated plant with EFNs is almost equally likely to live on islands as an abiotically pollinated plant without EFN, although we base this interpretation assuming strictly additive effects of EFNs and biotic pollination on island colonization, since we did not test for interactive effects of mutualistic traits (see more below). Similarly, in the island-level models, mutualism via EFNs positively predicts both ant and plant species richness on islands. Afkhami et al. (2014) proposed that defensive mutualisms, by protecting organisms from biotic interactions with negative fitness consequences, might facilitate range expansion, and Nathan et al. (2023) found evidence of this pattern in legumes that have EFNs and ant defenders. Establishment on an island involves both colonization and niche expansion, and generalist mutualists, such as in defensive ant-plant mutualisms mediated by EFNs, may be more likely to undergo both the expansion and ecological release phases of species establishment (Chomicki et al., 2019). Alternatively, rather than helping taxa colonize islands, species engaged in ant-plant mutualisms might be overrepresented on islands because they have larger population sizes and therefore reduced extinction rates (Weber and Agrawal, 2014).

Because there are few plant taxa with some combinations of mutualistic traits (e.g., very few plants have no mutualisms at all, see Figure 1), we did not explore the interactive effects of multiple mutualisms on insularity or island species richness. However, plants very commonly participate simultaneously or sequentially in multiple mutualisms, which can have non-additive ecological or evolutionary effects (Afkhami et al., 2014, Primieri et al., 2022, Laurich et al., 2023). For example, legumes with extrafloral nectaries that attract ants may also have some combination of rhizobia, mycorrhizae, and animal pollinators (Nathan et al., 2023). Even if biotic pollination constrains island colonization, the net result of multiple mutualisms on insularity might be positive, given that the positive effects of EFNs and mycorrhizae on insularity swamp the negative effect of biotic pollination in Table 2. However, multiple mutualisms could also result in non-additive effects on island species richness that are difficult to predict.

Mutualism can beget or hinder species diversity in ecological communities, with effects often depending on whether the benefits of mutualism are private to only one or a few community members, or widely available to many species (Bshary 2003, Frederickson et al., 2005, Rudgers and Clay, 2008). We examined the correlation between ant and plant species richness across oceanic islands, and found that the species richness of EFN-visiting ants was significantly predicted by the species richness of EFN-bearing plants, but not by plant diversity more generally. In contrast, the species richness of ants that do not visit EFNs was significantly predicted by non-EFN plant richness, but not the richness of EFN-bearing plants. These correlations suggest that ant and plant species that engage in mutualism via EFNs either respond more similarly to environmental variation across islands than, for example, EFN-visiting ants and non-EFN plant species, or that the establishment of one mutualist on an island accelerates the establishment of other mutualists in a process akin to invasional meltdown (Simberloff and Von Holle, 1999, Prior et al., 2015). However, the benefits of this kind of positive feedback appear restricted to plant and ant species that directly participate in the mutualism. In a global biogeographic analysis of ant-plant mutualisms, Luo et al. (2023) also found that the presence of potential ant partners was a driver of EFN-bearing plant species richness, suggesting that the pattern we observed might hold on both islands and mainlands.

Understanding what factors shape biodiversity on islands is critical because islands have unique biotas and island species are more vulnerable to extinction than mainland species (Losos and Ricklefs, 2009, Whittaker et al., 2017). We wholeheartedly agree with Delavaux et al. (2024) that mutualism plays an underappreciated role in island biogeography, but our findings suggest that mutualism does not universally filter island species richness, weakening the LDG. Instead, the facultative, generalized mutualism between ants and EFN-bearing plants is associated with increased insularity and island species richness. Furthermore, after accounting for phylogeny and other plant functional traits, only biotic pollination, and not mutualism with mycorrhizae or nitrogen-fixing symbionts, appears to reduce plant colonization of oceanic islands. Because mutualism links the fates of interacting species, mutualism makes key contributions to global biodiversity patterns, but most plant mutualisms do not prevent species from living on tropical islands as Delavaux et al. (2024) claimed.

## Authorship statement

DB and MEF designed the study, DB performed modeling work and analyzed data with input from MEF, DB wrote the first draft of the manuscript, and both authors jointly revised the manuscript for submission.

## Data statement

This paper uses publicly available data for analyses. All data sources are cited in the manuscript and code. All code is available on GitHub (https://github.com/drinnnird-uoft/island-mainland-EFN) and Dryad (reviewer URL http://datadryad.org/share/rvT89oxwB2Xu3pCoSUSf8GhHcxnHuFZl69ojStbSphE which will be replaced with a DOI url upon acceptance).

## Acknowledgements

We thank the Frederickson Lab for helpful feedback as this project developed. We also thank all the natural historians who have documented ant and plant trait data that culminated in the databases used in this study.

## Funding

M.E.F. thanks the NSERC Discovery Grant program for financial support.

## Novelty statement

Delavaux et al. (2024) claimed that mutualism filters plant diversity on islands, thereby weakening the latitudinal diversity gradient on islands. We re-analyzed their data, adding a new mutualism type: the defensive mutualism between plants with extrafloral nectaries (EFNs) and the ants that protect them from herbivores. By analyzing the geographic distributions of mutualistic plants and their ant partners on islands and mainlands, we found that ant and plant species that interact mutualistically via EFNs are over-represented on oceanic islands compared to other taxa. Thus, counter to Delavaux et al. (2024), this defensive ant-plant mutualism enhances rather than filters species richness on islands. Furthermore, this mutualism does not weaken the LDG on islands.

## Notes

### Competing Interest Statement

The authors have declared no competing interest.

### Summary of Updates

New data from Novais et al. (2025) on what ant species visit extrafloral nectaries were added to the analyses; additional models added to the Supplement.

