## Supplemental Material for "Defensive mutualisms affect plant and ant island biogeography at a global scale"

***Additional description of datasets***

The GIFT database is a set of geographic regions by latitude and longitude, such as islands or mainland locations, along with comprehensive species presence checklists for each region. Each region includes biogeographical variables and geological information. Each species checklist is a flat list of genus and species names that are known to be present in a particular geographic region. Trait data contained in GIFT includes information on 109 traits including morphology (woodiness, growth form, etc), life history (lifecycle, lifespan, etc), reproduction (selfing, seed measurements, etc), physiology (photosynthetic pathway, succulence, etc), genetics (ploidy, etc), and ecology (habitat, defense, etc). The trait data originates from a combination of 429 original floral checklists (see Wegelt et al. (2020) appendix) as well as using taxonomic and hierarchical trait derivations to assign traits at the genus/species level where appropriate. We used data on biotic pollination from GIFT. In addition, we used data on mycorrhizal associations at the species level from Soudzilovskaia et al. (2020), leveraging the species-level assignments where possible, otherwise assigning from the genus-level, and nitrogen fixer status from Werner et al. (2014), following Delavaux et al. (2024).

GABI records ant presence checklists by species, linked to spatially referenced polygons rather than latitude/longitude coordinates. GABI-I provides ant presence checklists on islands by latitude and longitude. The World List of Plants with Extrafloral Nectaries is a flat list of plant species that have EFNs. Similarly, the data from Kaur et al. (2019) is a flat list of ant species and whether or not they visit EFNs. The data from Novais et al. (2025) is a list of ant species and the plant taxa they visit, along with the locations of the EFNs. We converted this to a list of ant species that interact with EFNs. Each of these databases has been used in previous studies. They complement each other as they can all be tied to either species name or geographic location. Data availability for mainland sites was limited in some geographic regions, and many ant species described in GABI/-I had not yet been assigned scientific names, so they could not be matched with the Kaur et al. (2019) dataset or with the Novais et al. (2025) dataset. Oceanic islands have high endemism, where species present may not occur anywhere else, so there may be gaps in the trait data for these unknown plants and ants.

***Supplementary figures***

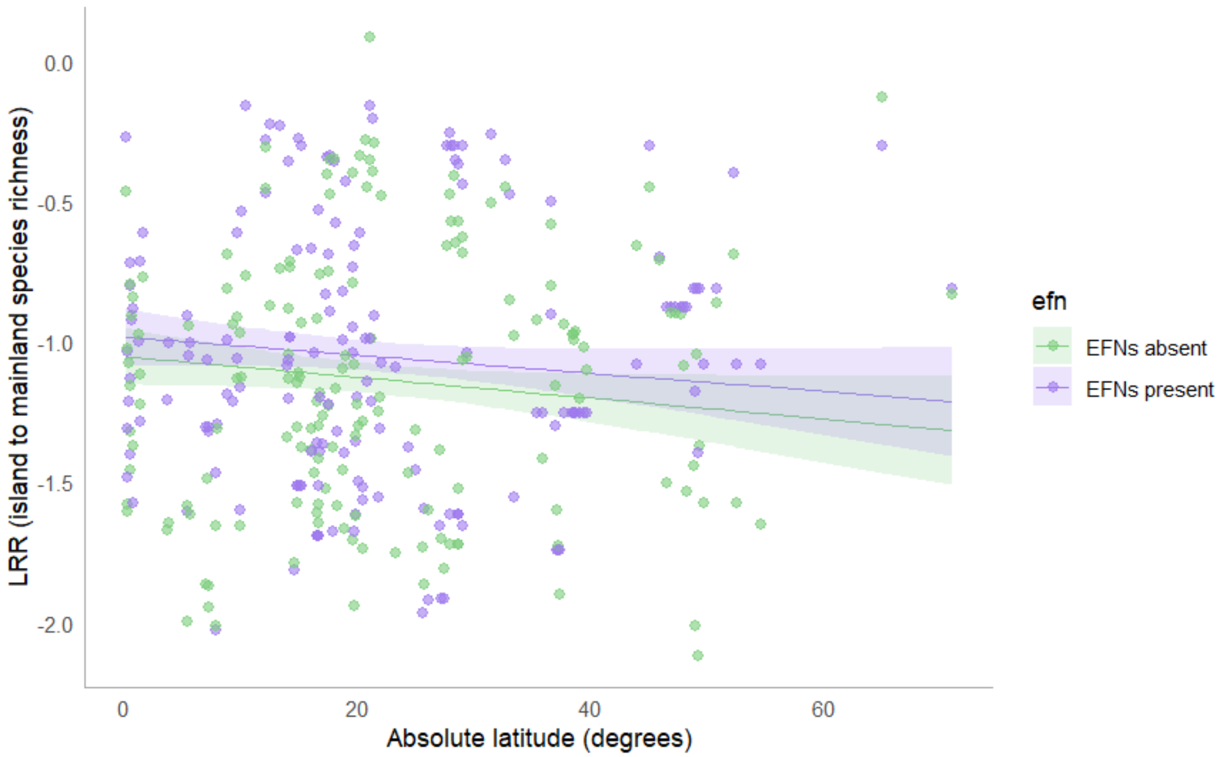


**Figure S1.** Plot of log response ratio of island to mainland species richness for EFN- (purple) and non-EFN-bearing (green) plants against absolute latitude of the island. Unlike in Figure 3, island-mainland species overlap was not required; no island species were excluded even if they did not occur on likely source mainlands.

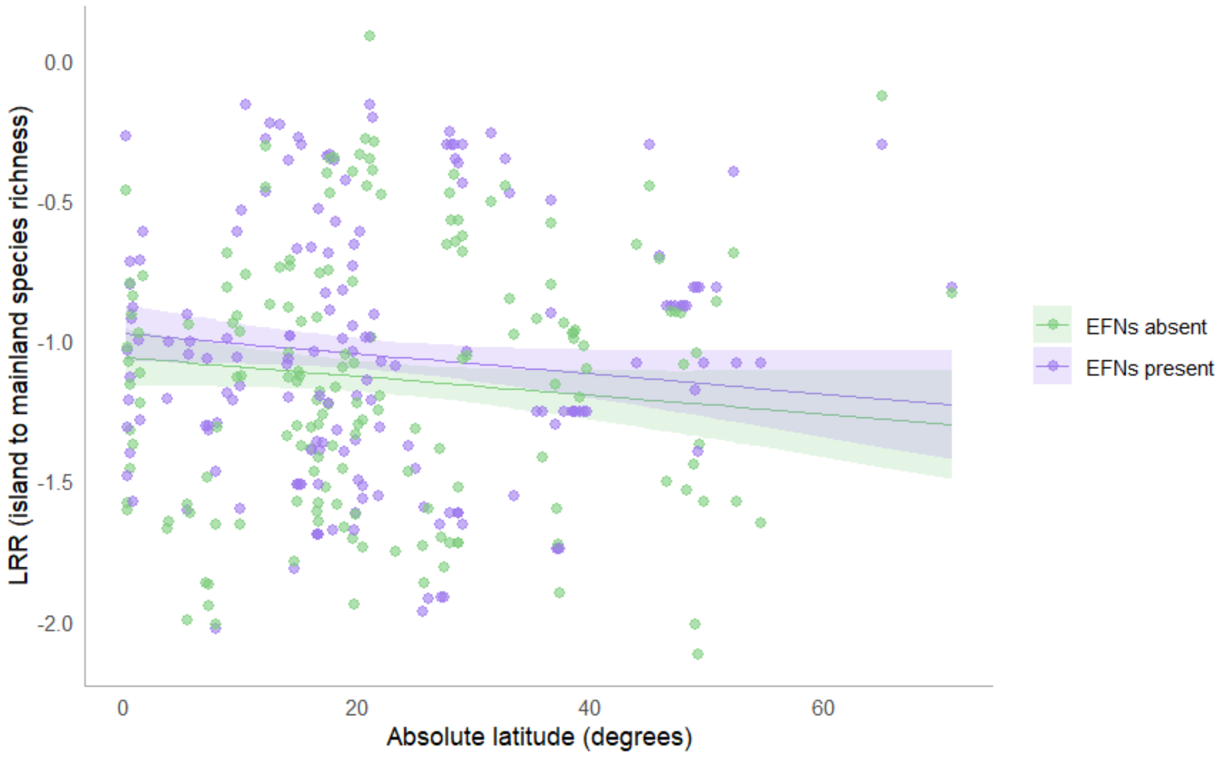


**Figure S2.** Plot of log response ratio of island to mainland species richness for EFN-visiting (purple) and non-EFN-visiting (green) ants against absolute latitude of the island. Unlike in Figure 4, island-mainland species overlap was not required; no island species were excluded even if they did not occur on likely source mainlands.

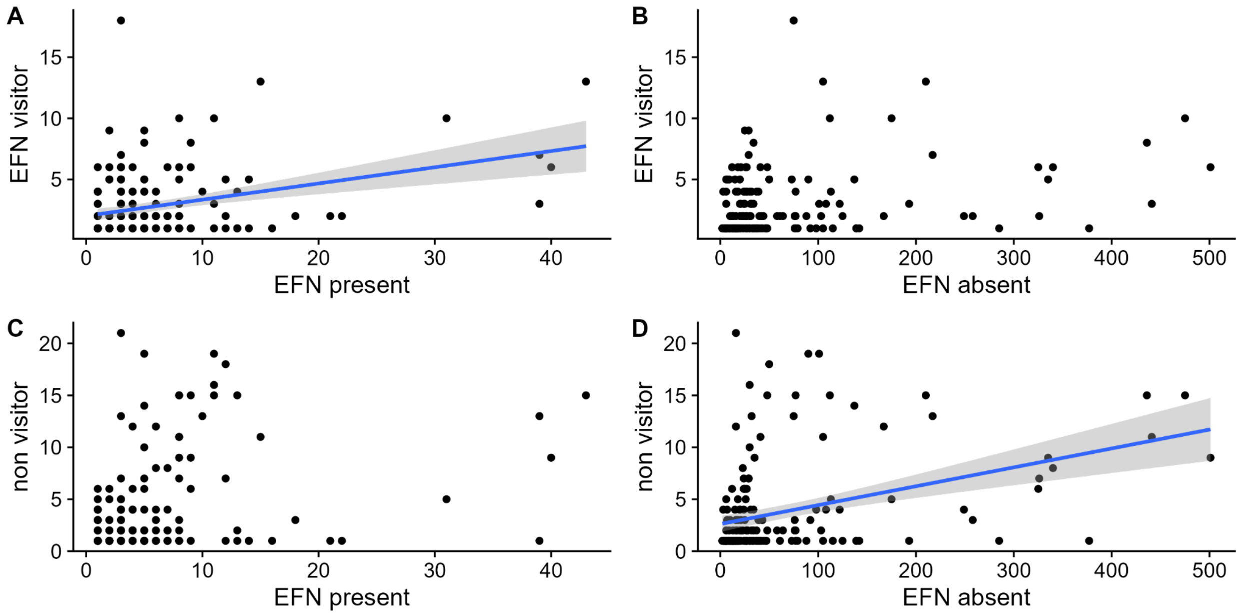


**Figure S3.** Correlations between ant and plant species richness across oceanic islands, according to mutualism status. A) Species richness of EFN-visiting ants (y-axis) against species richness of EFN-bearing plants (x-axis). B) Species richness of EFN-visiting ants (y-axis) against species richness of non-EFN plants (x-axis). C) Species richness of non-EFN-visiting ants (y-axis) against species richness of EFN-bearing plants (x-axis). D) Species richness of non-EFN-visiting ants (y-axis) against species richness of non-EFN plants (x-axis). Regression lines (blue) and confidence intervals (grey shaded regions) are shown for significant relationships only; see Table S2 for model results.

***Supplementary tables***

**Table S1**. Estimates and standard errors (s.e.) from Gaussian generalized linear models of the log response ratio of island to mainland species richness for A) plants (N = 169 groups) and B) ants (N = 178 groups) as a function of EFN mutualism status and biogeographical variables. Unlike in Table 3, island-mainland species overlap was not required; no island species were excluded even if they did not occur on likely source mainlands. Parameters ending in .i are island, .ml are mainland.

| | Estimate | s.e. |
|---|---|---|
| **A - Plants** | | |
| *Intercept* | *-1.06**** | *0.05* |
| *Absolute latitude.i* | -0.003 | 0.002 |
| *EFN?* | 0.09 | 0.07 |
| *Distance.ml* | *-0.15**** | *0.03* |
| *Area.i* | *0.25**** | *0.025* |
| *Area.ml* | *0.11** | *0.04* |
| Elevational range.i | 0.02 | 0.03 |
| Elevational range.ml | *0.08** | *0.03* |
| *Annual precip.i* | *0.21**** | *0.03* |
| Latitude * EFN | -0.0002 | 0.003 |

| **B - Ants** | | |
|---|---|---|
| *Intercept* | *-1.465**** | *0.101* |
| *Absolute latitude.i* | *-0.010*** | *0.004* |
| *EFN?* | *0.885**** | *0.108* |
| Distance.ml | 0.017 | 0.041 |
| *Area.i* | *0.157*** | *0.031* |
| *Area.ml* | *0.120** | *0.060* |
| Annual precip.i | -0.053 | 0.040 |
| Latitude * EFN | -0.007 | 0.004 |

Significant effects are *italicized* and indicated by *** for $p \leq 0.001$, ** for $p \leq 0.01$ and * for $p \leq 0.05$.

**Table S2.** Estimates and standard errors (s.e.) from Gaussian GLMs of EFN-visiting ant species richness as a function of the species richness of plants with and without EFNs. Parameters ending in .i are island, .ml are mainland.

| | Estimate | s.e |
|---|---|---|
| **A - Ants that visit EFNs** | | |
| Intercept | *3.90**** | *0.659* |
| *EFN-bearing plant species richness* | *0.067** | *0.034* |
| non-EFN bearing plant species richness | 0.002 | 0.003 |
| *Absolute latitude.i* | *-0.037*** | *0.014* |
| *Area.i* | *0.651*** | *0.202* |
| *Distance.ml* | *-3.150E-04** | *1.404E-04* |
| Annual precip.i | -0.268 | 0.236 |
| **B - Ants that do not visit EFNs** | | |
| *Intercept* | *4.629**** | *1.174* |
| EFN bearing plant species richness | 0.048 | 0.061 |

| | | |
|---|---|---|
| *non-EFN bearing plant species richness* | *0.014*** | *0.005* |
| Absolute latitude.i | -0.048 | 0.025 |
| Area.i | 0.595 | 0.360 |
| Distance.ml | -0.0005 | 0.0003 |
| *Annual precip.i* | *1.000** | *0.419* |

Significant effects are *italicized* and indicated by *** for $p \leq 0.001$, ** for $p \leq 0.01$ and * for $p \leq 0.05$.

**Table S3:** Estimates and standard errors (s.e.) from Gaussian generalized linear models of the log response ratio of island to mainland species richness for A) plants (N = 169 groups) and B) ants (N = 178 groups) as a function of EFN mutualism status and biogeographical variables. Unlike in Table 3, species were not filtered by GBIF occurrence. All species presence records were included. Parameters ending in .i are island, .ml are mainland.

| | Estimate | s.e. |
|---|---|---|
| **A - Plants** | | |
| *Intercept* | *-2.257**** | *0.06* |
| *Absolute latitude.i* | *0.006*** | *0.002* |
| *EFN?* | *0.833**** | *0.08* |
| *Distance.ml* | *-0.327**** | *0.03* |
| *Area.i* | *0.25**** | *0.03* |
| Area.ml | 0.049 | 0.05 |
| Elevational range.i | -0.012 | 0.03 |
| Elevational range.ml | 0.0002 | 0.04 |
| *Annual precip.i* | *0.23**** | *0.03* |
| *Latitude * EFN* | *-0.012**** | *0.003* |
| **B - Ants** | | |
| *Intercept* | *-2.335**** | *0.084* |

| | | |
|---|---|---|
| Absolute latitude.i | 0.003 | 0.003 |
| EFN? | 1.121*** | 0.090 |
| Distance.ml | -0.087* | 0.034 |
| Area.i | 0.100*** | 0.026 |
| Area.ml | 0.003 | 0.050 |
| Annual precip.i | -0.053 | 0.033 |
| Latitude * EFN | -0.008** | 0.004 |
